# NAMPT activity plays a key role in driving autoimmune processes that characterise type 1 diabetes development in mice

**DOI:** 10.1101/2025.01.14.632985

**Authors:** Daniel Egbase, Sophie R. Sayers, Naila Haq, Jithu Varghese, Vesela Gesheva, Sreya Bhattacharya, Jay Kynaston, Ella L. Hubber, Lorna Smith, Timothy J. Pullen, Henry Gerdes, Vivian K. Lee, David Hopkins, Min Zhao, Yee Cheah, Joshua Greally, Sam Butterworth, James A Pearson, Gavin Bewick, Shanta Persaud, Paul W. Caton

## Abstract

Type 1 diabetes (T1D) is characterised by destruction of pancreatic beta cells by islet-infiltrating cytotoxic lymphocytes, and elevated intra-islet secretion of pro-inflammatory cytokines. However, the underlying pathophysiological mechanisms remain incompletely understood. We hypothesised that abnormal elevation of islet NAD, via activation of NAMPT, plays a key role in driving islet autoimmune processes in T1D.

Here, we report that NAMPT inhibition protects against pro-inflammatory cytokine (IL-1β, TNFα and IFNγ) mediated beta-cell dysfunction and apoptosis in isolated mouse and human islets. RNAseq revealed that NAMPT inhibition blocked cytokine-mediated gene expression linked to pro-inflammatory responses and leukocyte migration. In vivo, diabetes was induced in CD1 mice via multiple low dose streptozotocin (MLDS) injection. MLDS mice were administered the NAMPT inhibitor FK866 (10 mg/kg; IP) or saline equivalent for 16 days. These experiments demonstrated that NAMPT inhibition improved glycaemic control and beta-cell function and insulin content in MLDS mice. FK866 also reduced proportions of islet-residing TNFα-producing CD4^+^T-cells and F4/80^+^macrophages, proliferation of spleen-derived CD4^+^ and CD8^+^T-cells, and proliferation of islet-derived CD4^+^T-cells and F4/80^+^macrophages. Finally, we report that NAMPT inhibition was able to block pro-inflammatory cytokine-mediated migration of cytotoxic CD8^+^T-cells into isolated islets, using an in vitro transwell platform.

This data supports a key immunomodulatory role for NAMPT in islet autoimmunity. NAMPT inhibition may represent a novel therapeutic approach for T1D. The effects of increased NAD levels on islet inflammation require in-depth characterisation, and caution should be exercised with regard to use of NAD boosting supplements, particularly in individuals at risk of developing T1D.

## Introduction

Type 1 diabetes (T1D) is an autoimmune disorder characterised by auto-reactive immune cell-mediated destruction of insulin-producing pancreatic beta cells (1–4). The precise triggers that initiate pathogenesis of T1D remain unclear, with both genetic susceptibility and/or environmental triggers reported to prime autoimmune events in affected individuals (5–9). Autoimmune events in T1D are characterised by islet infiltration by CD4^+^ and CD8^+^T-cells, in a process known as insulitis (1–4, 10). Cytotoxic CD4^+^ and CD8^+^T-cells secrete pro-inflammatory cytokines (such as IL-1β, TNFα and IFNγ) within the intra-islet environment (11–13), resulting in autoimmune destruction of beta-cells.

Several issues surround the current standard-of-care for T1D, including hypoglycaemia associated with exogenous insulin therapy and poor graft survival following islet transplantation (14–22). Teplizumab, an anti-CD3 mAb which can delay development of T1D, was recently approved (23). However, no drugs are currently available which can prevent T1D.

One candidate as a regulator of islet inflammation and T1D drug target is nicotinamide phosphoribosyltransferase (NAMPT). NAMPT is well characterised as the rate-limiting enzyme in the NAD salvage pathway, where it catalyses the conversion of nicotinamide (NAM) to nicotinamide mononucleotide (NMN). NMN is subsequently converted to NAD by NMN adenynyltransferase enzymes (24). NAD+ metabolism regulates homeostasis of the innate and adaptive immune response (25). However, accumulating evidence indicates that NAMPT activity, potentially mediated via enhanced NAD+ levels, can facilitate dysregulation of immunomodulatory processes and onset of inflammation.

For example, NAMPT activity reportedly induces TNFα production in mouse and human inflammatory cells (16) and can indirectly induce T-cell activation and IFNγ production in mature T-cells (27–30). These effects, as well as enhanced T-cell proliferation are reduced in response to NAMPT inhibition. Interestingly, NAMPT inhibition has no effect on inactive T lymphocytes (31). NAMPT activity is also required for activation of pro-inflammatory M1 macrophages, and NAMPT inhibition downregulates TNFα expression in M1 macrophages and polarizes macrophages towards an anti-inflammatory M2 phenotype (32–34). Consistent with these findings, a key role for NAMPT activity in pathogenesis of auto-immune conditions including Inflammatory Bowel Disease, arthritis and autoimmune encephalomyelitis has been reported (31, 35). Further support for a role for NAD in mediating inflammation is derived from recent studies reporting that high levels of the NAD boosting precursors niacin and nicotinamide riboside (NR) can promote inflammation in macrophages and endothelial cells (36,37).

Crucially, many of the immunomodulatory effects described above are also key pathogenic events in T1D development. This suggests a potential role for NAMPT activity in the pathophysiology of islet autoimmunity and beta-cell destruction.

To investigate the therapeutic effects of NAMPT inhibition in T1D, we examined the anti-diabetic effects of two separate NAMPT inhibitors, FK866 (38,39) and compound 17 (C17; (40)), in the multiple-low dose streptozotocin (MLDS) T1D model (41) and in isolated mouse and human islets treated with pro-inflammatory cytokines.

## Research Design and Methods

### MIN6 cell culture

MIN6 cells were cultured at 37°C in DMEM (25 mmol/l glucose; 2 mmol/l glutamine, 10% FBS (vol./vol.), 100 U/ml penicillin, 100 μg/ml streptomycin) and treated with FK866 or C17 prior to NAD and NMN measurements.

### NMN and NAD+ Measurement

NMN was measured in a cell-free environment or in MIN6 β-cells using a previously described fluorometric assay (42,43). NAD levels were measured by an NAD/NADH Quantification Kit, (Sigma-Aldrich). See Supplementary methods.

### Pancreatic islet isolation and treatment

Mouse islets were isolated from 8-week-old non-diabetic CD1 mice, as previously described (44). Pancreas were inflated with 1mg/mL collagenase solution (Sigma-Aldrich, Poole, U.K.) followed by density gradient separation (Histopaque-1077; Sigma-Aldrich). Human islets were isolated from heart-beating non-diabetic donors (44) at the King’s College Hospital Human Islet Isolation Unit, with the appropriate ethical approval (KCL HI-RTB; 20/SW/0074). Isolated islets were maintained in supplemented RPMI at 37 °C and 5% CO_2_ in a humidified atmosphere for up to 24 hours prior to experimental treatment. For insulin secretion and apoptosis measurements ∼60 mouse islets were incubated with pro-inflammatory cytokines (0.05 U/ml Il-1β, 1 U/ml TNFα and IFNγ) +/- either FK866 (10 – 40 nM) or C17 (50 – 400 nM), or with FK866 (10 nM)/C17 (200nM) alone, for 24 h.

### Glucose-stimulated insulin secretion

Mouse CD1 or human islets were pre-incubated in physiological salt solution containing 2mM glucose. Groups of 5 size-matched islets were then further incubated at 37°C for 1 h in salt solution and 2 or 20 mmol/l glucose. Secreted insulin was measured using an in-house I125 radioimmunoassay (45).

### Islet apoptosis

Apoptosis was determined using a Caspase-Glo 3/7 luminescent assay (Promega, Southampton, UK).

### Bulk RNA Sequencing Analysis

Total RNA was extracted from ∼150 islets per mouse (RNeasy Mini Kit, Qiagen, UK). Total RNA concentration and purity was quantified by NanoDrop (ND-1000, Thermo Fisher Scientific, Hemel Hempstead, UK) spectrophotometer. RNAseq libraries were prepared from 1000 ng total RNA using a NEBNext Ultra II Directional RNA Library Prep kit (New England Biolabs, Hertfordshire, UK. Cat.no: E7760). Briefly, mRNA was isolated using the Poly-A mRNA magnetic isolation module (E7490) (New England Biolabs, Hertfordshire, UK). Isolated mRNA was enzymatically fragmented prior to cDNA synthesis. Each sample was then tagged with a single index that was unique to the sample. cDNA library was amplified by PCR for 12 cycles. The resulting library was purified using Agencourt AMpure XP beads (Beckman Coulter, Buckinghamshire, UK). Library quality was assessed using a Bioanlyzer 2100 and DNA High Sensitivity kit (Agilent). Library fragment size distribution was determined using Agilent 2100 expert software. Library concentration was determined by qPCR using NEBNext Library Quant Kit for Illumina (New England Biolabs (NEB), Hertfordshire, UK. Cat.no: E7630). Libraries were pooled at equal molar ratio and sequenced using an Illumina NovaSeq 6000 at 150bp paired end with an approximated depth of 25 million reads per sample.

### Diabetic mouse models

Diabetes was induced by multiple-low dose streptozotocin (MLDS) administration (41). 4–6-week-old CD1 mice (26 – 33g) (Charles River, UK) were randomly assigned into four groups: non-diabetic control (citric acid buffer (CAB)^Saline^), diabetic control (MLDS^Saline^), diabetic FK866-treated (MLDS^FK866^), and diabetic vehicle-treated (5% in DMSO in saline; MLDS^Vehicle^). To induce diabetes, mice were administered 40 mg/kg/body weight STZ (IP; Bio-techne, Bristol, UK; dissolved in CAB, pH 4.5) or CAB alone, for 5 consecutive days (days 1 – 5). For drug administration, FK866 (10 mg/kg body weight; IP) or vehicle control were administered daily (16 days; Days 7-22). Mice were housed in 12 h light/dark cycle, temperature-controlled conditions with *ad libitum* access to standard mouse chow and water. Procedures were performed in accordance with UK Home Office regulations (Animal Scientific Procedures Act, 1986).

### Diabetes evaluation

Ambient blood glucose levels were measured every 2 – 3 days at 10 am, and diabetes was defined by blood glucose levels >13.8 mM over 3 consecutive days. Blood samples were taken from the tail tip. Capillary blood glucose levels (mM) were determined using an Accucheck glucose meter (Roche Diagnostics, UK).

### Serum Insulin and C-peptide

Overnight fasted mice were administered 2 g/kg body weight 30% glucose solution (IP). After 30 min, mice were culled, and blood was collected. Plasma insulin (Mercodia, Sweden) and C-peptide (ALPCO, United States) were measured by specific ELISA.

### Pancreas histology

Whole pancreas was excised and fixed in 4% paraformaldehyde for 24 h prior to tissue processing (Leica TP 1020, Leica Biosystems, Germany) and paraffin embedding. 5 μm paraffin sections were melted briefly on a hot plate before the slides were dewaxed using xylene and then rehydrated using solutions of decreasing ethanol gradient (100%–70%). The staining procedure was as follows: 5 min immersion in haematoxylin, 5 min in running water, three to five dips in acid alcohol, 2 min in running water, five dips in eosin and then 2 further minutes in running water. After staining, the sections were dehydrated in an increasing ethanol gradient (70%–100%). Finally, the sections were immersed in xylene for 7 min before being mounted. Slides were imaged with a bright field channel on a microscope (Olympus BX40). A minimum of 60 islets per mouse (three mice/group) were scored for insulitis. Histological analysis of immune cell infiltrates in pancreatic islets was performed blinded by an independent observer (JK).

### Islet dissociation prior to flow cytometric analysis

Islets were isolated from MLDS mice, washed in PBS and dissociated with TrypLE Express (1 ml/100 islets) for 10 min at 37 °C, triturating every 5 min for 10 s. After washing with DMEM (supplemented with 10% FBS), then PBS, samples were transferred to round bottom tubes and kept on ice.

### Spleen dissociation and treatment prior to flow cytometric analysis

MLDS mouse splenocytes were isolated as previously described (46). On the day of sacrifice, spleens were removed, placed on ice in Hanks’ Balanced Salt Solution and mechanically disrupted using forceps. The cell suspensions were centrifuged at 1500 rpm (4°C) for 5 minutes and erythrocytes were lysed by suspension of the pellets in 5 ml of ACK (Ammonium-Chloride-Potassium) Lysing Buffer (ThermoFisher Scientific, Altrincham, UK). Cell suspensions were then incubated at room temperature for 10 minutes, followed by centrifugation. Supernatants were discarded, and pellets were washed twice with HBSS and resuspended in supplemented RPMI. An aliquot of cell suspension was mixed with 0.4% trypan blue solution (Sigma-Aldrich). Through the trypan blue exclusion method, and using a haemocytometer chamber the cells were counted. 1 × 106 cells/ml of cells were seeded on P24 round-bottom plates and cultured (37 °C 5% CO_2_) in 10 µM EdU for 24 h. To detach from the flask, cells were then incubated in a solution of trypsin/EDTA (0.1%/0.02%) for 2 min in an incubator at 37 °C, prior to addition of DMEM. After centrifugation at 1000 rpm for 2 min, the spleen cell pellet was re-suspended in fresh DMEM and kept on ice for flow cytofluorimetry analysis.

### Flow cytometric analysis

Single-cell suspensions were incubated with anti-mouse CD4, CD8 and F4/80 (all 1:50; to assess immune cell number), anti-mouse IFNγ or TNFα (both 1:50; immune cell activation) and anti-mouse insulin (1:100; beta-cell number) fluorochrome-conjugated monoclonal antibodies. Click-iT EdU Flow Cytometry Cell Proliferation Assay (Thermo Fisher Scientific, Hemel Hempstead, UK) was used to assess immune cell proliferation. See Supplementary Methods. Gating strategies are shown in Supplementary Figs S3-S9.

### Migration assays

Transwell CD8^+^T-cell migration assays were performed as previously described (47), using reformed islets originally isolated from BALB/C mice and haplotype-matched CD8+ T-cells isolated from 8-week-old diabetic female NY8.3 NOD mice. Cells were exposed to pro-inflammatory cytokines +/-FK866. After 18 h incubation, CD8^+^ cell migration into islets was quantified by measuring the O.D values at Ex/Em = 540/590 nm using a PHERAstar Microplate Reader (33). See Supplementary Methods. Figure 6 shows a schematic representation of the experimental set-up.

### Data analysis

Significance was tested using one- or two-way ANOVA with Tukey’s hoc test, using GraphPad PRISM software (San Diego, CA, USA). Data are expressed as mean ± SEM.

## Results

### NAMPT inhibition protects pancreatic islets from pro-inflammatory cytokine-induced dysfunction and apoptosis

Beta-cell dysfunction and death in T1D is partly mediated by chronic exposure to pro-inflammatory cytokines. To determine the importance of NAMPT in beta-cell failure in T1D we exposed mouse and human islets to pro-inflammatory cytokines with or without NAMPT inhibitors (FK866 or C17; Supplementary Fig.1A-G). Static glucose stimulated insulin secretion (GSIS) studies showed that both FK886 and C17 exerted significant protective effects against cytokine-mediated reductions in GSIS (both p=<0.0001 vs. Cytokines 1 ng/ml) (Fig. 1A and 1B). Neither FK866 (10 – 20 nM) nor C17 had any effects on static GSIS when added in the absence of cytokines (Supplementary Fig. 2A). Co-incubation and pre-treatment with FK866 (Fig. 1C and Fig. 1D; p=<0.0001 vs. Cytokines 1 ng/ml) or compound 17 (Fig. 1E) also significantly protected against apoptosis (caspase 3/7 activity) in mouse islets. Finally, we observed that both FK866 and C17 also prevented cytokine-mediate apoptosis in human islets (Fig.1F). FK866 had no effect on caspase 3/7 activity in the absence of cytokines (Supplementary Fig. 2B).

**Fig. 1.**
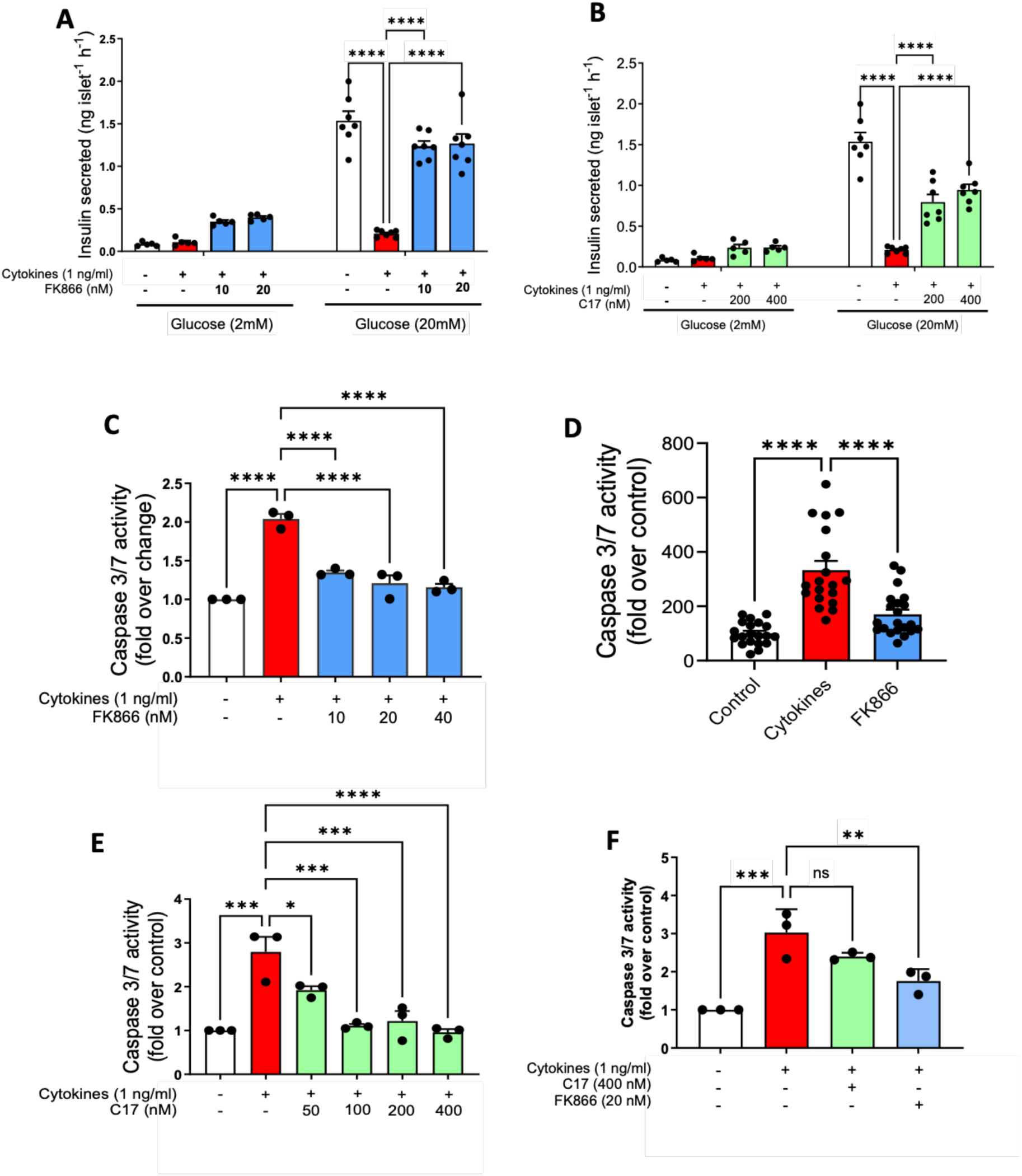
NAMPT inhibition exerts protective effects against pro-inflammatory cytokine-induced beta-cell dysfunction and apoptosis in isolated pancreatic islets. **(A-B)** Static insulin secretion in response to basal (2mM) or 20mM glucose was assessed in isolated mouse islets incubated with cytokines (CK; IL-1β, TNFα and IFNγ; 1 ng/ml), with or without either (A) FK866 (10 – 20 nM) or (B) compound 17 (200 – 400 nM) for 24 h. N = 5 – 7 replicates from 6 mice and representative of three independent experiments. (**C-D**) Apoptosis (caspase 3/7 activity) was measured in isolated mouse islets treated with (C) CK cocktail with or without FK866 (10 – 40 nM) for 24 h, or (**D**) in mouse islets pre-treated with FK866 (10 nM) for 2 h followed by CK treatment (without FK866) for 22 h. (**E**) Apoptosis (caspase 3/7 activity) measured in isolated mouse islets treated with CK cocktail with or without compound 17 (50 – 400 nM) for 24 h. For C-E, n = 3 where an n of 1 is equal one experiment with 10 wells, each with five size-matched islets. (**F**) Apoptosis (caspase 3/7 activity) measured in isolated human islets treated with a CK cocktail with or without either FK866 (20 nM) or compound 17 (400 nM) for 24 h. n = 3 where an n of 1 is equal one experiment with 10 wells, each with five size-matched islets. Values in all graphs are represented as means ± SEM. Statistical significance between groups is depicted by *p < 0.05, **p<0.01; **p<0.001 ****p < 0.0001.

### FK866 protects pancreatic mouse islets from deleterious pro-inflammatory cytokine-mediated transcriptional changes

To understand how NAMPT activity impacts the deleterious effects of pro-inflammatory cytokines in greater detail, we conducted transcriptome analysis on isolated mouse islets co-treated with cytokines and FK866 via bulk RNA sequencing (RNA-seq). Cytokine treatment led to the significant upregulation of 1873 genes and downregulation of 1718 genes (Fig. 2A). Gene set enrichment analysis (GSEA) revealed most elevated responses were downstream of cytokine activity (IFNγ signalling and TNF*α*+ signalling). Key beta cell functional and maturity genes were also downregulated (Fig. 2B and 2C). More in-depth analysis of the “Inflammatory response” gene sets illustrates many of the cytokine-induced significantly upregulated genes encode for proteins mediating or responding to inflammation and cellular stress. These include chemokines (*Ccl5*, *Cxcl10*, *Cxcl9*, *Ccl22*), cytokine receptor proteins (*Tnfsf10*, *Tnfrsf1b*, *Osmr*) and antigen processing and presenting proteins (*Tapbp, Cd40, Marco*) likely symbolising islet cells experiencing a pro-inflammatory response (Fig. 2D and 2E).

**Fig. 2.**
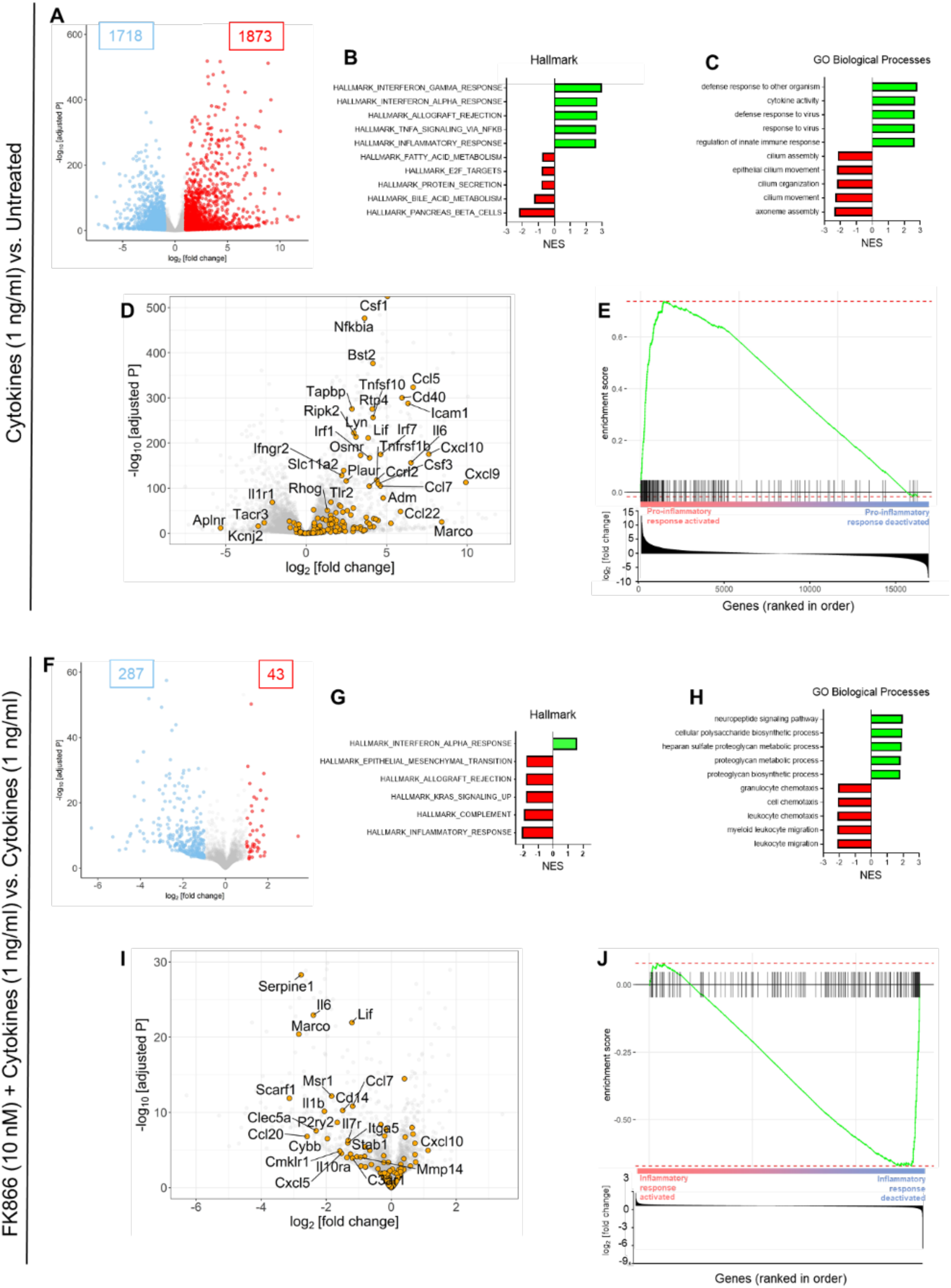
FK866 suppresses cytokine-induced inflammatory transcriptional pathway enrichment in isolated pancreatic mouse islets. **(A and F)** Volcano plot depicting log2 fold changes of differentially expressed (DE) annotated genes from RNA-seq performed on isolated mouse islets treated for 24 h with **(A)** a cytokine cocktail (IL-1β, TNFα and IFNγ; 1 ng/ml) versus untreated, or **(F)** FK866 (10 nM) and a cytokine cocktail versus a cytokine cocktail. Each dot represents a gene. DE genes with a log2 fold change of >1, all adjusted P < 0.05 are highlighted in red (indicating upregulation) or blue (indicating downregulation). Numbers denote total number of upregulated and downregulated genes between treatment groups. **(B,C,G,H)** Five most and least enriched pathways as revealed by gene set enrichment analysis (GSEA) in isolated mouse islets treated for 24 h with **(B and C)** a CK cocktail versus untreated, or **(G and H)** FK866 (10 nM) and a cytokine cocktail versus a cytokine cocktail. GSEA performed with **(C,G)** ‘Hallmark’ and **(B,H)** ‘Gene Ontology (GO) biological processes’ gene sets from the Molecular Signatures Database (MSigDB). Only gene sets with significant (p < 0.05) normalised enrichment scores shown. **(D and I)** Volcano plot depicting log2 fold changes of DE genes within the ‘HALLMARK_INFLAMMATORY_RESPONSE’ gene set. The annotated genes are from RNA-seq performed on isolated mouse islets treated for 24 h with **(D)** a CK cocktail versus untreated, or **(I)** FK866 (10 nM) and a cytokine cocktail versus a cytokine cocktail. Each dot represents a gene. Gene names are displayed for top differentially expressed genes (log2 fold change >1, adjusted P < 0.05). **(E and J)** Enrichment plots derived from the GSEA performed on the transcripts within the ‘HALLMARK_INFLAMMATORY_RESPONSE’ gene set in isolated mouse islets treated for 24 h with **(E)** a CK cocktail versus untreated, or **(J)** FK866 (10 nM) and a cytokine cocktail versus a cytokine cocktail. Green line denotes running statistic and black lines represent a single gene in the gene set. Bottom left figures are bar plots showing all DE genes log2 fold changes ranked in order. n=3 independent biological sample replicates (each replicate consisted of 150 islets).

Compared to cytokine-treated islets, FK866 + cytokine treated islets had 43 and 287 upregulated and downregulated genes, respectively (Fig. 2F). Notably, the presence of FK866 in the treatment led islet cells to downregulate pathways related to leukocyte migration, as well as leukocyte and granulocyte chemotaxis. Interestingly, FK866 reversed cytokine-induced upregulation of the “Inflammatory response” gene set (Fig. 2G and 2H). Specifically, FK866 reversed the cytokine-induced expression of chemokines and chemokine receptors (Ccl20, Cxcl5, Cmklr1), cytokine receptor proteins (*Il10r*, *Il7r*, *Osmr*), and cytokines (*Il1b*, *Il6* and *Lif*), innate immune response genes (*Clec5a*, *Cybb*) (Fig. 2I). Leading edge analysis also exemplifies that FK866 supressed a pro-inflammatory response in islets (Fig. 2J).

### NAMPT inhibition ameliorates hyperglycaemia and reductions in insulin content in MLDS-induced diabetic mice

Given that NAMPT activity played a role in cytokine-induced beta-cell function, survival and transcriptomics *in vitro*, we next sought to assess the effects of pharmacological NAMPT inhibition with FK866 on the course of diabetes development *in vivo*. Diabetes was induced in mice by multiple-low dose streptozotocin (MLDS) administration (40 mg/kg bw; IP for 5 consecutive days). FK866 was administered to MLDS-induced diabetic mice, beginning at day 7 (two days after the end of STZ administration) as outlined in Fig. 3A. From day 11 until the end of the experiment (day 23), FK866 treatment significantly lowered (∼25%) ambient blood glucose concentrations in MLDS^FK866^ mice, compared to MLDS^Saline^ and MLDS^Vehicle^ treatment (Fig. 3B and 3C). FK866 administration also lowered fasting blood glucose levels in MLDS mice (Fig. 3D).

**Fig. 3.**
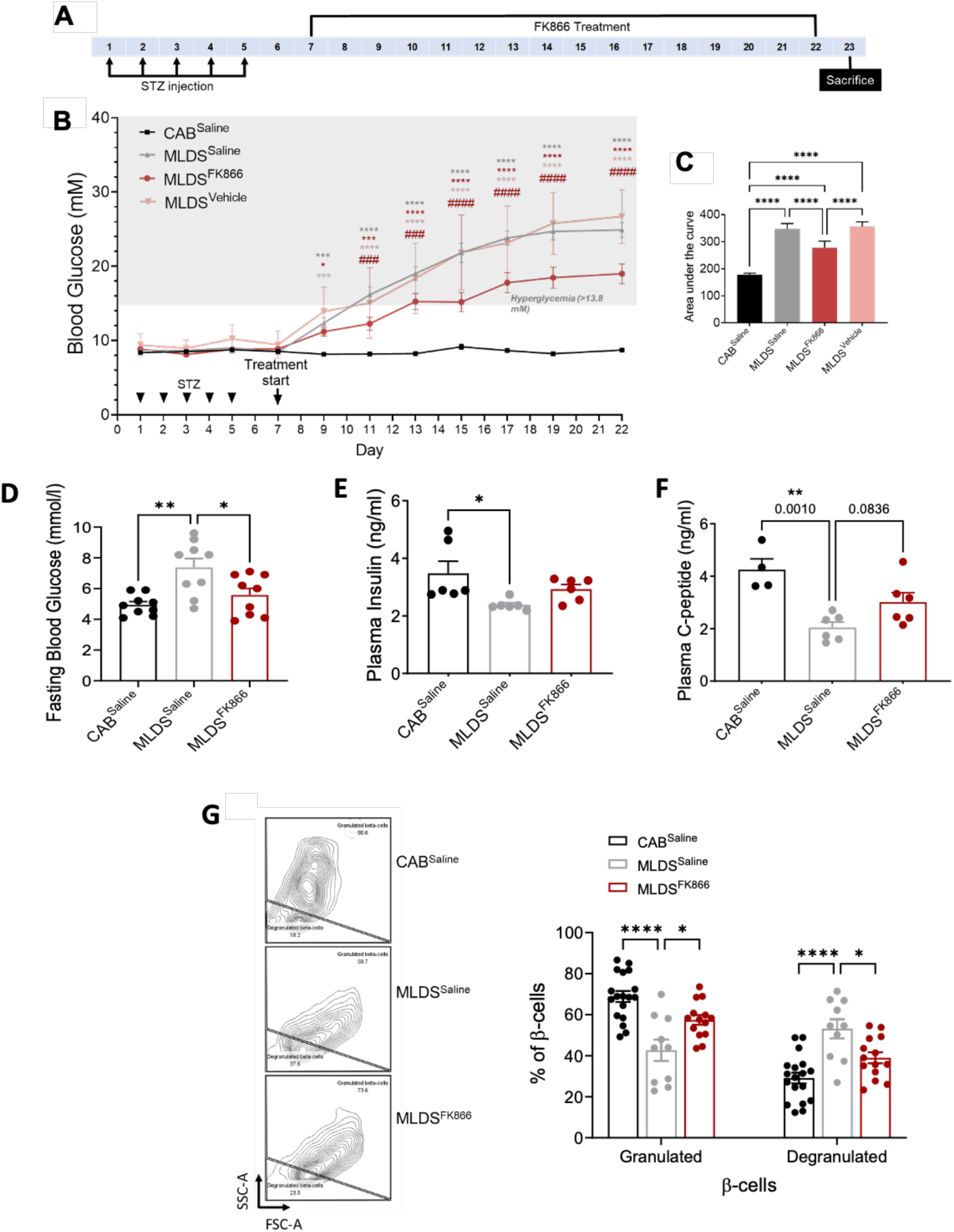
FK866 ameliorates MLDS-induced hyperglycaemia and reduced beta-cell functionality. **(A)** Schematic diagram of the experimental protocol: mice were administered citric acid buffer (CAB) or multiple low dose streptozotocin (MLDS) regimen, then after 48 hours, were administered saline, FK866 or vehicle (IP) for 17 days. **(B and C)** Ambient blood glucose measured every 2-3 days and the corresponding area under the curve analysis. In **(B)**, black arrows and annotations denote streptozotocin (STZ) exposure and FK866 treatment (n = 12-29). **(D)** Fasting blood glucose (day 23); **(E and F)** Plasma levels of insulin and C-peptide measured by ELISA using terminal blood samples taken on the day of sacrifice (day 23) following fasting and then injection of 2 g/kg 30% glucose solution (n = 6). **(G)** Representative image of data showing SSC/FCS of granulated and degranulated β cells on the day of sacrifice (day 23). Also shown are corresponding percentages of the optical properties of insulin-positive islet cells following isolation, dispersion and immunofluorescence staining of insulin. n = 14-18 pools of islet cells of which a maximum of 5000 cells were analysed. Values in all graphs are represented as means ± SEM. Statistical significance vs CAB^Saline^ is depicted by *p < 0.05, **p < 0.01, ***p < 0.001, ****p < 0.0001. Statistical significance vs MLDS^Saline^ is depicted by #p < 0.05, ##p < 0.01, ###p < 0.001, ####p < 0.0001.

To determine whether improvements in glycaemic controls are related to improvements in beta-cell function, we assessed whether FK866 enhanced serum insulin and c-peptide response to glucose. MLDS^Saline^ mice exhibited significantly lower plasma insulin (p=0.05) and C-peptide (p=0.01) compared to non-diabetic/CAB^saline^ animals. FK866 treatment slightly, but non-significantly, prevented MLDS-mediated reductions in insulin (Fig. 3E) and c-peptide secretion (3F, p<0.08). Beta cell health was examined in greater detail through assessment of insulin content per beta cell, using flow cytometry. MLDS induced a significant reduction in sub-population of granulated beta cells (p=0.0001) and a significant increase in sub-population of degranulated beta cells (p=0.0001). Consistent with a beta-cell protective effect, FK866 treatment led to a significant increase in proportion of granulated beta-cells (p<0.05) and a significant decrease in proportion of degranulated beta-cells (p<0.05) compared to the MLDS^Saline^ (Fig. 3G). These findings suggest that pharmacological NAMPT inhibition with FK866 improves glycaemic control, in part through partial protection of beta-cell function and maintenance of beta-cell insulin content during MLDS-induced diabetes development.

### NAMPT inhibition protects MLDS-induced diabetic mice from insulitis

We next sought to determine whether the effects of FK866 on glycaemic control were linked to supressed insulitis. Whilst all severities of insulitis were observed in MLDS^Saline^ and MLDS^FK866^ mice, MLDS^FK866^ mice exhibited a greater proportion of islets that remained in the perivascular/periductal infiltration stage rather than progressing to the peri-insulitis/insulitis phase (Fig. 4A - C), indicating partial protection against insulitis by FK866. To gain a deeper insight into these effects, the proportion of immune cell sub-types present within the islet was examined using flow cytometry. MLDS^Saline^ mice displayed a ∼2-fold increase of CD4^+^T-cells (p=0.05), CD8^+^T-cells (p<0.01), and F4/80^+^macrophages (p<0.001), in the islet cell population compared to non-diabetic controls (Fig. 4D – F). Similar to the outcomes observed following H+E staining, FK866 administration led to a partial decrease (∼25%) in the proportion of CD4^+^T-cells, CD8^+^T-cells and F4/80^+^macrophages in the islet population, compared to MLDS^Saline^ (Fig. 4D - F). A crucial factor in immune-mediated T1D pathophysiology is activation of immune cells leading to TNFα and IFNγ production which in turn mediate beta-cell destruction. To assess the effect of FK866 on immune cell activation, we began by measuring TNF*α* and IFNγ production within the intra-islet environment. MLDS^Saline^ mice had significantly elevated proportions of TNF*α*^+^ cells in the total islet population compared to non-diabetic mice (p=0.05), and FK866 blocked this effect (Fig. 4G; P<0.01). Assessment of immune sub-populations within all TNF*α*^+^ cells revealed MLDS^Saline^ mice displayed significantly increased proportions of TNF*α*^+^CD4+ (P<0.05) and TNF*α*^+^F4/80^+^ cells (P<0.0001) compared to non-diabetic control mice, and these increases were blocked by FK866 (p=0.01 and p=0.001, respectively; Fig. 4H and 4I). MLDS^Saline^ mice also displayed significantly elevated proportions of IFNγ^+^ cells in the total islet population (p<0.05; Supplementary Fig. Fig. 10A), and partial increases in IFNγ^+^CD4+ and IFNγ ^+^F4/80^+^ sub-populations (Supplemental Fig. 10B -C). FK866 administration led to small but non-significant reductions in these effects. Together this data demonstrates that NAMPT inhibition can prevent islet immune-cell activation, specifically by reducing TNFα production from islet CD4^+^T-cells and F4/80^+^macrophages. This provides a plausible mechanism through which FK866 prevents beta-cell destruction and improves glycaemic control in MLDS mice.

**Fig. 4.**
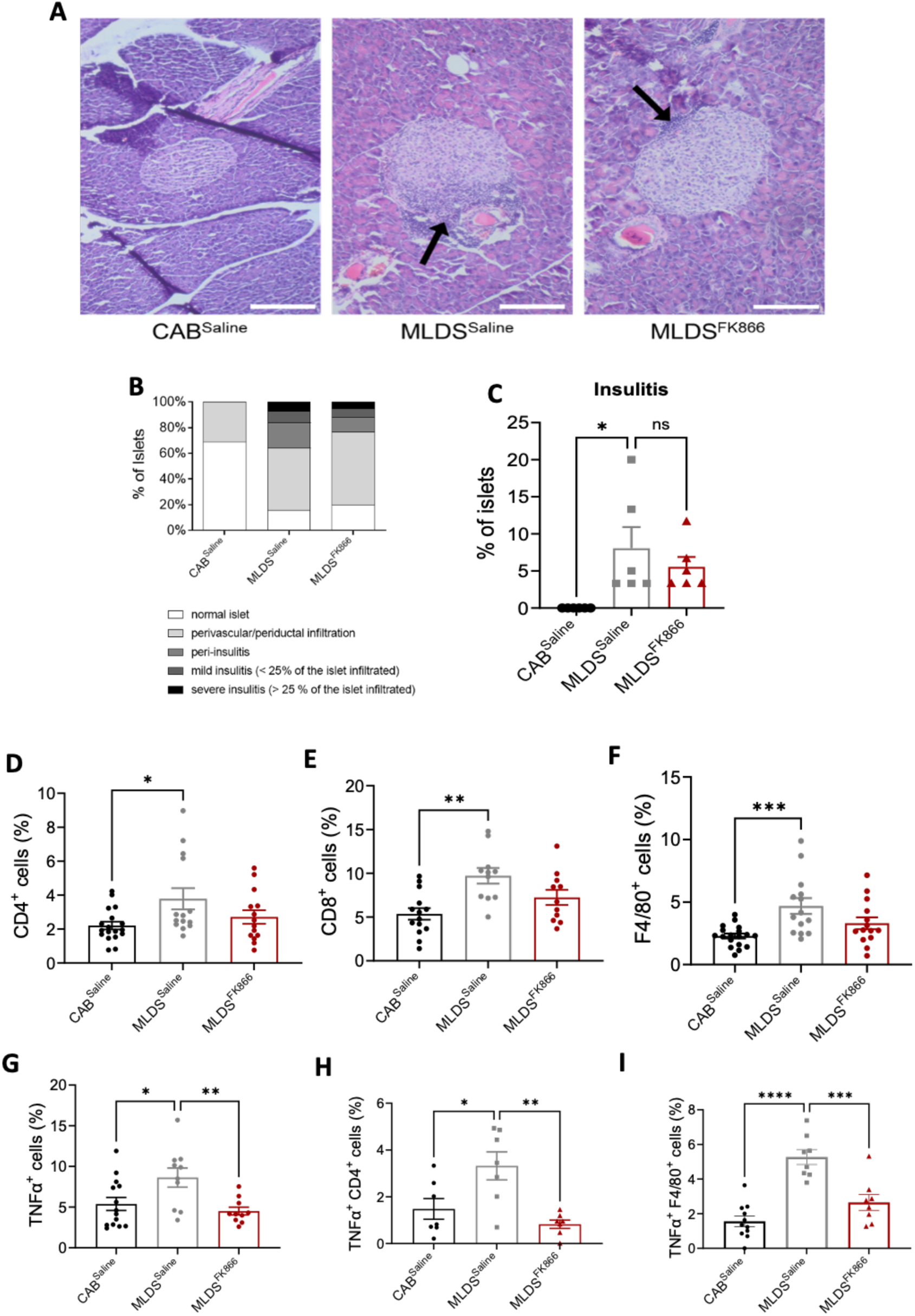
FK866 partially protects MLDS-induced diabetic mice from insulitis. **(A)** Representative images of mouse pancreas from CAB^Saline^, MLDS^Saline^ and MLDS^FK866^ mice stained with hematoxylin and eosin and evaluated for severity of infiltrating immune cells using light microscopy. Black arrows indicate immune cell infiltration. Imaged at 20× (Scale bars, 50 μM). **(B)** Corresponding percentages of islets graded for all stages of insulitis. **(C)** Percentages of islets showing mild and severe insulitis only. Data generated from 60 islets per mouse pancreas (three mice per group). **(D-F)** Percentage of **(D)** CD4^+^, CD8^+^ and **(F)** F4/80^+^ cells within the islet cell population of CAB^Saline^, MLDS^Saline^ and MLDS^FK866^ mice following sacrifice. n = 14-18 pools of islets of which a maximum of 5000 cells were analysed. **(G-I)** Percentages of **(G)** TNFα+, **(H)** TNFα+CD4+, **(I)** TNFα+F4/80+, within the islet cell of CAB^Saline^, MLDS^Saline^ and MLDS^FK866^ mice. n = 7-14 pools of islets of which a maximum of 5000 cells were analysed. Values in all graphs are represented as means ± SEM. Statistical significance is depicted by *p < 0.05, **p < 0.01, ***p < 0.001, ****p<0.0001.

### NAMPT inhibition reduces splenic- and islet-derived immune cell proliferation in MLDS-induced diabetic mice

To further elucidate the immunomodulatory impact of NAMPT inhibition, we examined the effects of FK866 on proliferation of islet-derived CD4^+^ and CD8^+^ T-cells and F4/80^+^ macrophages. MLDS^Saline^ mice displayed significantly increased proportions of proliferating CD4^+^, CD8^+^ and F4/80^+^ cells compared to non-diabetic mice (p<0.01, p<0.05 and p<0.05, respectively) (Fig. 5A-C).FK866 prevented MLDS-induced increases in islet-derived CD4^+^T-cells and F4/80^+^ cell proliferation (Fig. 5A and 5B; p<0.05), but not CD8^+^T-cells proliferation (Fig. 5C). FK866 also partially prevented MLDS-induced increases (p<0.05) in spontaneous splenocyte proliferation (Fig. 5D). Staining for immune cell sub-types within the splenocyte population revealed that FK866 also prevented MLDS-mediated increases in spontaneous proliferation of spleen-derived CD4^+^ and CD8^+^T-cells (P<0.05), (Fig. 5E-F). Together, these results demonstrate the clear anti-inflammatory, immunomodulatory effect of NAMPT inhibition on CD4^+^ and F4/80^+^ immune cells activation during T1D development.

**Fig. 5.**
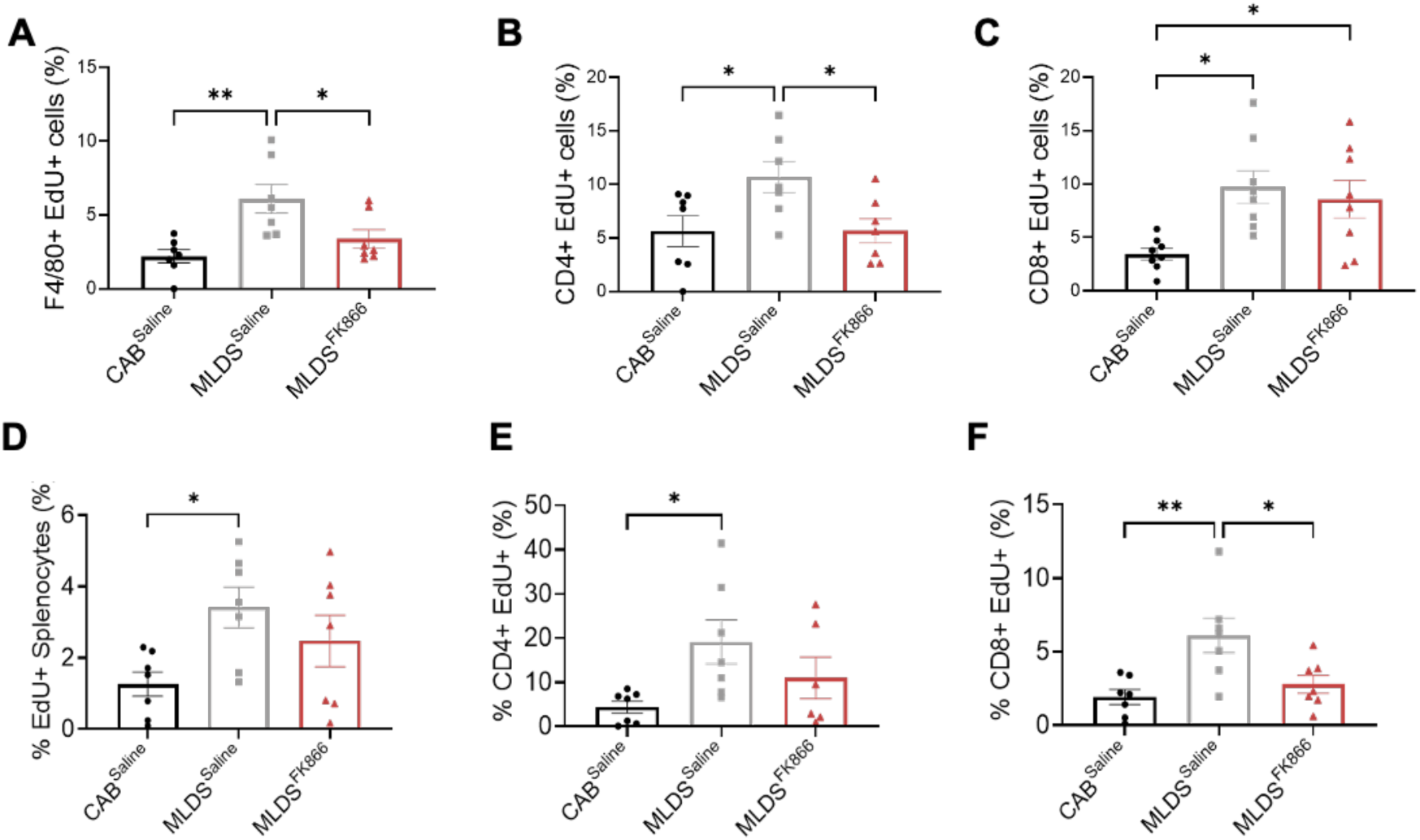
FK866 blocks MLDS-mediated increased proliferation of islet- and spleen-derived immune cells. **(A-C)** Percentages of **(A)** F4/80^+^EdU^+^, **(B)** CD4^+^EdU^+^, **(C)** CD8^+^EdU^+^ cells within the islet cell population of CAB^Saline^, MLDS^Saline^ and MLDS^FK866^ mice following sacrifice, pancreatic islet isolation, dispersion, EdU-supplemented culture (72 h) staining and flow cytometric analysis. **(D-F)** Percentages of **(D)** EdU^+^ splenocytes, **(E)** CD4^+^EdU^+^, **(F)** CD8^+^EdU^+^ cells within the splenocyte population of CAB^Saline^, MLDS^Saline^ and MLDS^FK866^ mice following sacrifice, spleen removal, dispersion, culture with EdU (24 h), staining and flow cytometric analysis. n = 7-8 pools of islet or spleen cells of which a maximum of 5000 cells were analysed. Values in all graphs are represented as means ± SEM. Statistical significance is depicted by *p < 0.05, **p < 0.01.

### NAMPT inhibition prevents cytotoxic CD8^+^T-cell migration into pancreatic islets

To further examine the immunoregulatory effects of FK866, we utilised a transwell culture platform to assess the effect of FK866 on migration of CD8+T-cells obtained from NOD mice, into pancreatic islets (Fig. 6A). Exposure of islets to pro-inflammatory cytokines alone led to a 3-fold increase in CD8+T-cell migration compared to untreated islets (p<0.0001). However, when islets were either pre-treated with FK866 prior to cytokine exposure, or co-incubated with FK866 and cytokines, these effects were completely blocked, with CD8+T-cell migration remaining at basal levels (p<0.0001) (Fig.6B).

**Fig. 6.**
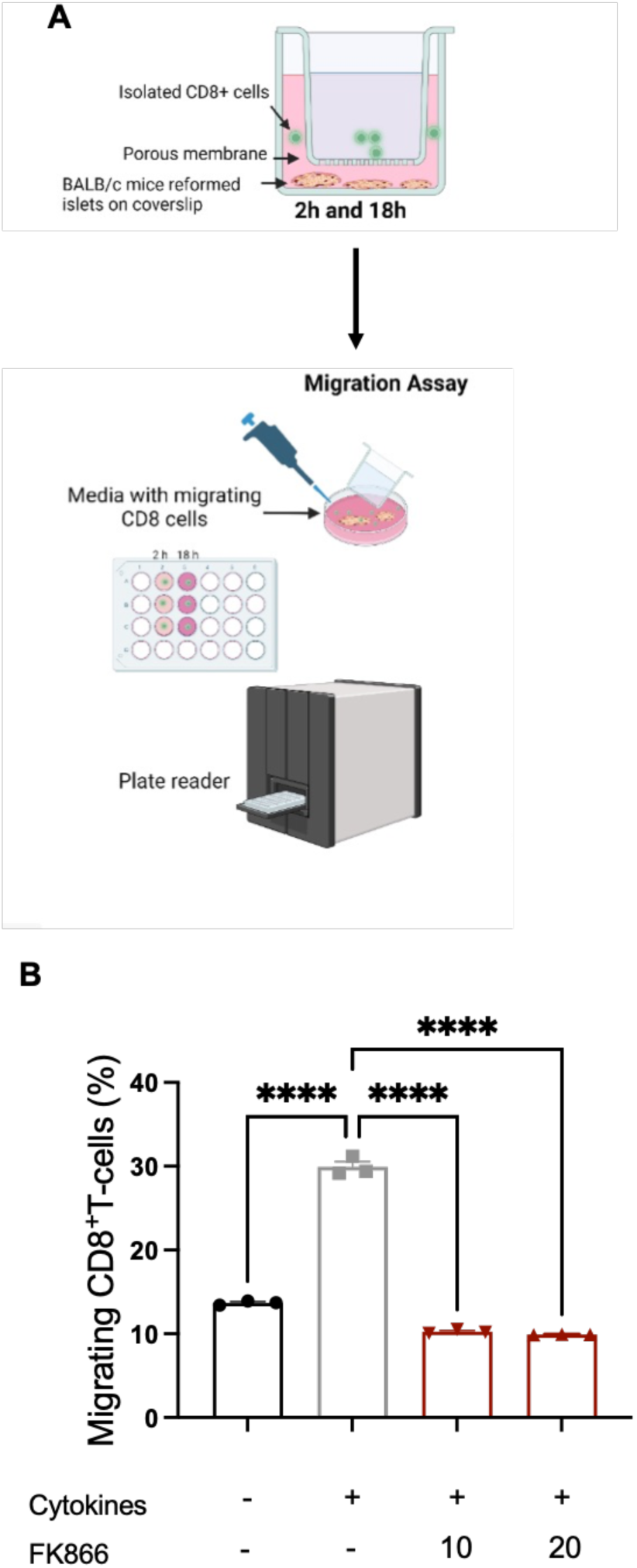
FK866 prevents migration of cytotoxic CD8^+^T-cells into pancreatic islets. **(A)** Schematic diagram of the experimental protocol. Islets were isolated from BALB/c mice and dispersed and reaggregated into reformed islets. CD8+T-cells were isolated from splenocytes derived from NY8.3 NOD mice. **(B)** Islets were exposed to pro-inflammatory cytokines (IL-1β, TNFα and IFNγ; 1 ng/ml), with or without FK866 (10 – 20 nM) for 18 h. Migration of CD8+ cells into islets was assessed using a transwell culture platform. Values in all graphs are represented as means ± SEM. Statistical significance is depicted by ****p < 0.0001.

## Discussion

Here, we provide evidence that supports a role for NAMPT as an immunomodulatory enzyme with a key role in T1D pathogenesis. Of potential clinical importance, NAMPT inhibition leads to improved glycaemic control and beta-cell function in MLDS-induced diabetic mice.

Clinical studies report that T1D patients with stimulated plasma C-peptide levels of as little as ≥0.02 pmol/mL (indicative of residual beta-cell function) exhibit improved metabolic control and lower insulin requirements, compared to T1D patients with plasma C-peptide levels below this threshold (48,49). This suggests that relatively small improvements in beta-cell function, as observed in this study, can lead to clinically relevant improvements in glycemic control in T1D.

These beneficial effects are driven in part by prevention of insulitis and islet immune cell activation. In T1D, auto-immune beta-cell destruction is attributed predominately to increased islet levels of IL1β, TNFα and IFNγ (50–52). NAMPT inhibition *in vivo* reduced MLDS-induced IFNγ and TNFα cytokine production in islet cells. This is in line with previous reports demonstrating that dysregulated NAD+ metabolism is essential for CD4^+^T-cell IFNγ and TNFα secretion (53). Furthermore, we demonstrated that FK866 can prevent migration of NOD mouse derived CD8^+^T-cells into pancreatic islets. Cytotoxic CD8^+^T-cells are key drivers of T1D pathophysiology through mediation of beta-cell destruction and the ability of FK866 to block migration of these cells into islets suggests therapeutic utility of this compound.

Transcriptional analysis suggested that NAMPT activity plays a key role in cytokine-induced leukocyte migration and chemotaxis and TNFA- and IFNG-related pathways, providing a potential explanation of the prohibitive effects on insulitis observed in response to NAMPT inhibition *in vivo*. In addition to transcriptional changes, FK866 may also impact the secretion of peptides that stimulate macrophage and leukocyte recruitment (54). NAMPT exists in intra-cellular (iNAMPT) and extra-cellular (eNAMPT) forms. FK866 has been shown to block IFNγ- and TNFα-induced eNAMPT secretion from keratinocytes and fibroblasts. Moreover, eNAMPT sensitises TNFα-induced expression and secretion of CCL2, CXCL8, and IL6 which induce immune cell recruitment (55) and play a key role in insulitis (56–59). We report that MLDS-induced diabetes development led to significantly increased islet IFNγ and TNFα levels. This suggests that MLDS-induced islet inflammation could mediate increased eNAMPT release within the local islet environment, which promotes insulitis via increased expression of CCL2, CXCL8, and/or IL6. Furthermore, FK866 treatment may block eNAMPT release and subsequent expression of immune cell recruitment molecules, which will reduce insulitis. Further investigations into eNAMPT release from islet cells, and its role in the immunopathology of T1D are required.

Elevated spontaneous proliferation of islet-derived T-cells and macrophages have been characterised as part of MLDS-induced diabetes pathophysiology (32). These effects were confirmed in our investigations and were blocked by NAMPT inhibition. This is consistent with studies reporting FK866-mediated inhibition of T-cell proliferation and survival *in vitro* (46). FK866 can also induce T-cell dysfunction by depleting intracellular ATP levels via impeding glycolysis (60). NAD can also be metabolised by CD38 and CD157 to generate cADPR and NAADP, which can activate Ca^2+^ signalling pathways critical for T-cell proliferation (61). Thus, FK866 may prevent increased CD4^+^T-cell proliferation through indirectly reducing Ca^2+^ availability. Similarly, high levels of the NAD precursors niacin and NR have been reported to promote inflammation via activation of CD38 and generation of metabolites 2PY and 4PY (36, 37), and FK866 may impact islet inflammation by blocking these pathways.

Our data showed that FK866 protected against cytokine-induced reductions in GSIS in isolated mouse islets. It should be noted that a previous study found that NMN administration, potentially via increased NAD synthesis, protected isolated mouse islets from cytokine-attenuated GSIS (62–65). However, these studies typically utilised levels of NMN (∼100 μM) several orders of magnitude greater than are observed physiologically. Therefore, it is difficult to compare the findings of these experiments with the current study, in which islet NMN levels were generally <10 μM across all treatment groups.

We report that NAMPT inhibition protects isolated mouse and human islets from cytokine-induced apoptosis. At first glance, these results are surprising as treating cancer cells with FK866 induces apoptosis following increased caspases-3/-7 activation (66). However, a recent study found that FK866 supressed caspase-3 activity in H_2_O_2_ stimulated-HeLa cells, but increased LDH release (a measure of necrosis) (67). The reasoning behind these findings is that as NAMPT activity regulates NAD+, and subsequently ATP levels; and because ATP fuels caspase-3 activation, NAMPT therefore plays a key role in determining whether cells undergo caspase-3-mediated apoptosis or caspase-3-independent necrosis. For pancreatic beta cells, changes in cytokine signalling, NAD+ salvage, and intracellular ATP levels could independently or concurrently factor into whether they survive or die. Thus, there may be an ideal, pharmacological level of NAMPT inhibition in pancreatic beta cells that prevents of extrinsic apoptosis pathway yet does not trigger necrosis. Further investigations are necessary to elucidate the mechanisms linking intracellular NAD+ metabolism and beta cell survival.

## Conclusion

In summary, we characterise NAMPT as an immunoregulatory enzyme that plays a key role in autoimmune events observed in T1D development in mice. The data presented here demonstrate proof-of-concept identifying NAMPT inhibition as a novel therapeutic intervention to prevent autoimmune beta-cell destruction during T1D development. Moreover, further research is required to determine the effects of increased NAD levels on islet inflammation, and caution should be exercised with regard to use of NAD boosting supplements, particularly in individuals at risk of developing type 1 diabetes.

## Acknowledgements

We thank J. I. Miyazaki (Osaka University, Osaka, Japan) for provision and consent to use MIN6 cells. Flow cytometric analysis was conducted in the Biomedical Research Centre Advanced Flow Cytometry Platform at King’s College London (London, UK). Library generated for RNA sequencing was conducted by the Genomics Centre at King’s College London. Human islets were provided through the King’s College London Human Islet Research Tissue Bank (KCL HI-RTB; 20/SW/0074). We thank the families of the pancreas donors and the Cell Therapy Unit in the NIHR/Wellcome King’s Clinical Research Facility at that houses the clinical islet isolation laboratory for making islets available for this research. D.E. was recipient of a King’s Medical Research Trust PhD studentship. This work was also supported by an EFSD/Lilly European Diabetes Research Programme grant and Diabetes UK project grants (18/0005865; 20/0006297). The study funders were not involved in the design of the study; the collection, analysis, and interpretation of data; writing the report; or the decision to submit the report for publication.

## Author contributions

DE, SRS, NH, JJV, VG, SB, VKL, ELH, LS, HG, JK, TJP, KJP, DH, MZ, SB, JG, JP, GB, SJP and PWC designed and conducted research, analysed data, and reviewed and approved the final manuscript. DE and PWC wrote the paper. PWC is the guarantor of this work.

