## Supplementary Methods and Data for "NAMPT activity plays a key role in driving autoimmune processes that characterise type 1 diabetes development in mice"

**NMN and NAD+ Measurement**

For cell-free experiments, reaction mixtures were prepared in triplicate containing the following: 1ml of Reaction Buffer (made up of PRPP (Sigma, UK), 2 mM ATP, 0.02% bovine serum albumin (BSA), 2 mM dithiothreitol (DTT), 12 mM MgCl2, and 50 mM Tris–HCl (pH 7.5)), 2ug/ml recombinant NAMPT and FK866 (5 – 20 nM) or compound 17 (50 – 200 nM). For NMN measurements in cells, MIN6 were maintained in monolayer culture. Then, 20,000 MIN6 β-cells were extracted with 100 µl of 1 mol/L perchloric acid and then neutralized by addition of 330 mL of 3 mol/L K2CO3. This was followed by incubation at 4 °C for 10 min, and centrifugation at 12000 x g for 15 min at 4 °C, with the supernatant retained. 50 ml of samples (reaction mixtures or cellular samples) and of standard solutions of NMN (50 ml; 25 – 200 mmol/l) were subsequently derivatised by addition of 20 ml of 1 mol/L KOH and 20 ml acetophenone (Sigma, Poole, UK) followed by incubation at 4°C for 15 min. 90 ml Formic acid was then added, and the solution incubated for 10 min at 37°C, producing a highly fluorescent compound. 150 ml of samples or standards were then added into a 96-well plate fluorescence was detected on a SpectraMax i3x plate reader with excitation and emission wavelength of 382 and 445 nm, respectively. NAD was measured by specific colorimetric assays, according to manufacturer's instructions (Sigma-Aldrich).

Preparation of reformed islets

2000-2500 islets isolated from BALB/c mice (Envigo, UK) were hand-picked from suspension cultures 24h after isolation and washed with PBS. Islets were then dissociated into a suspension of single cells through trypsin digestion. Trypsin digestion was stopped by adding neurobasal media consisting of MEM supplemented with 5% (v/v) FBS, 1x B-27, 1%(v/v) penicillin and streptomycin, HEPES 10 mM, Glutamax 1x, Na-pyruvate 1mM, and 11 mM glucose. Islet cells were seeded at a density of 35,000 cells/cm$^2$ on laminin-coated or collagenase-coated glass coverslips. Cells were then cultured for 3-4 days to allow adherence and spread on the glass surfaces, and for a further 10-14 days to form reformed islets.

**Migration Assays**

Islets were isolated from BALB/c mice, and reformed islets were generated as described above. BALB/c mice express the MHC class H2Kd haplotype and are therefore matched to antigen-specific T cells. CD8+ cells were isolated from the spleens of 8-week-old diabetic female NY8.3 NOD mice (a kind gift from Dr James Pearson, Cardiff University). A single-cell suspension was prepared with MojoSort buffer at 1 × $10^8$ cells/ml. 10 µl of biotinylated antibody containing antibodies against CD4, CD11b, CD11c, CD19, CD24, CD45R/B220, CD49b, CD105, I-A/I-E, TER-119/Erythroid, and TCRγδ were added. The mixture was then incubated for 15 min on ice, and 10 µL streptavidin nanobeads were added. The final volume of 120 µL of the cell and bead mixture was incubated on ice for an additional 15 min. Finally, 120 µL was diluted in in 1x MojoSort buffer (2.5 mL) and placed in a MojoSort Magnet (Cat# 480019/480020, BioLegend) for 5 min for magnetic separation. The free cells in the solution were collected as enriched $CD8^+$ T cells.

The migration assay was conducted using an 8µm pore transwell plate (BioVision, Inc., USA) according to the manufacturer's protocol. BALB/c islets were incubated for 48h in 2% FBS RPMI complete medium in the lower chamber of a Transwell culture plate. Following this, 2X$10^5$ $CD8^+$ T cells were seeded into the upper chamber and placed over coverslips with reformed islets treated with pro-inflammatory alone or in combination with FK866. After 18 h of incubation, the cells in the lower chamber were quantified by measuring the O.D values at Ex/Em = 540/590 nm using a PHERAstar Microplate Reader (33). Figure 6 shows a schematic representation of the experimental set-up.

**Flow cytometric analysis**

To assess viability, single-cell suspensions were stained with LIVE/DEAD™ Fixable Blue Dead Cell Stain (Thermo Fisher Scientific). To assess surface immune cell markers, single-cell suspensions were incubated with anti-mouse CD4, CD8 and F4/80 fluorochrome-conjugated monoclonal antibodies (all 1:50 dilution), or with isotype-matched controls (Becton Dickinson, Wokingham, UK) for 45 minutes at RT. Following staining, cells were fixed and permeabilized with Intracellular Fixation & Permeabilization Buffer Set (Thermo Fisher Scientific, UK). To assess for proliferative cells, Click-iT EdU Flow Cytometry Cell Proliferation Assay (Thermo Fisher Scientific, Hemel Hempstead, UK) was used according to manufacturer's instructions. Briefly, cells were resuspended in 100 µL of 1X Click-iT® saponin-based permeabilization, then 0.5 mL of Click-iT® reaction cocktail was added to each sample for 30 minutes at room temperature, protected from light. Samples were washed twice with 1 mL of 1X Click-iT® saponin-based permeabilization and wash reagent, followed by resuspension in eBioscience™ Flow Cytometry Staining Buffer (Thermo Fisher Scientific, UK). To assess other intracellular proteins, single-cell suspensions were incubated with anti-mouse insulin (1:100), IFNγ or TNFα (both 1:50) fluorochrome-conjugated monoclonal antibodies or with isotype-matched controls (Becton Dickinson, Wokingham, UK) for 45 min at RT. 5000 islet cells/pool or 100000 spleen cells/spleen were analysed on a BD LSRFortessa™ using BD FACSDiva™ software (BD Biosciences, Southampton, UK) and analysis was conducted with FlowJo (Tree Star). Gating strategies are shown in Supplemental Figure 3-9.

### Supplementary Figures

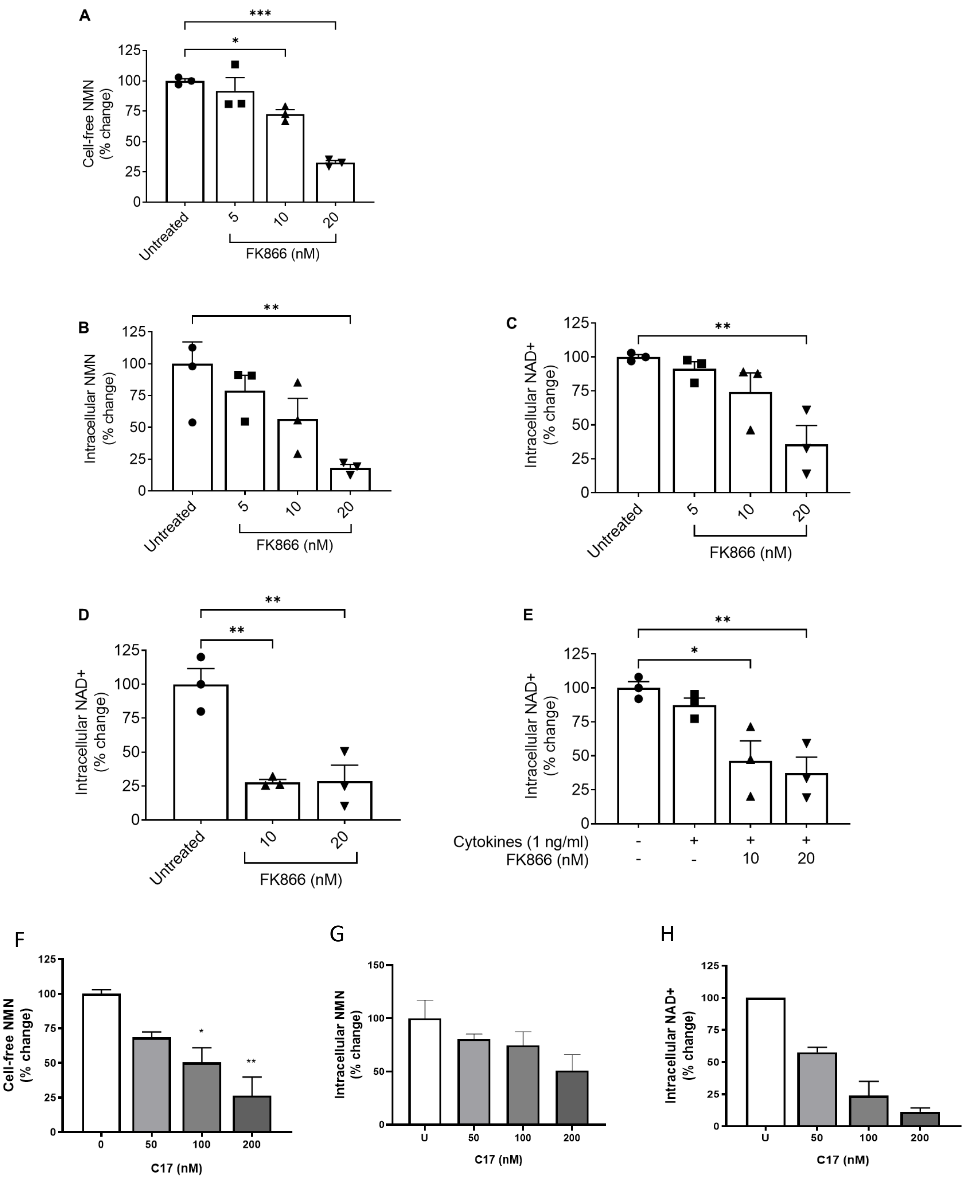


**Supplementary Figure 1. NAMPT inhibition *by* FK866 *and Compound 17.* (A)** NMN levels detected in reaction mixtures containing reaction buffer, wild-type recombinant human-NAMPT, NAM, FK866 (5 – 20nM) were incubated at 37°C for 15 min. n=3. **(B)** NMN levels detected in MIN6 beta-cells cultured for 24 hr with FK866 (5 – 20nM). **(C)** NAD+ levels measured in MIN6 beta-cells cultured for 24 hr with FK866 (5 – 20nM). **(D)** NAD+ levels measured in isolated mouse pancreatic islets cultured for 24 hr with FK866 (10 and 20 nM). **(E)** NAD+ levels measured in isolated mouse pancreatic islets cultured for 24 hr with a cocktail of proinflammatory cytokines (IL-1β, TNFα and IFNγ; 1 ng/ml), or with both a cytokine cocktail and FK866 (10 and 20 nM). **(*F*)** NMN levels detected in reaction mixtures containing reaction buffer, wild-type recombinant human-NAMPT, NAM, compound 17 (5*0* – 20*0* nM) were incubated at 37°C for 15 min. n=3. **(*G*)** NMN levels detected in MIN6 beta-cells cultured for 24 hr with FK866 (5 – 20nM). **(*H*)** NAD+ levels measured in MIN6 beta-cells cultured for 24 hr with FK866 (5 – 20nM). Values in all graphs are represented as means ± SEM. Statistical significance between groups is depicted by *p < 0.05, **p<0.01; **p<0.001.

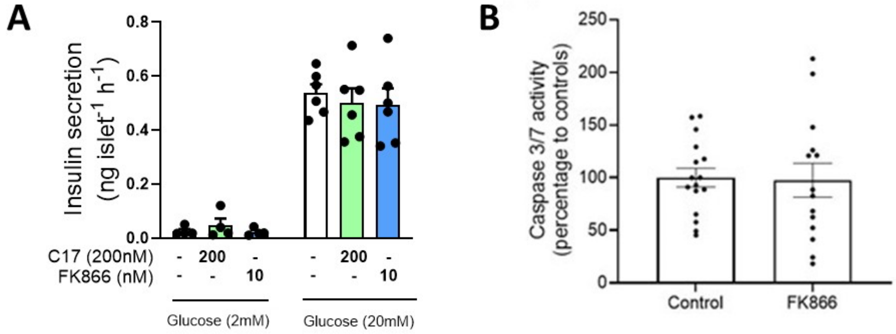


**Supplementary Fig. 2. NAMPT inhibition has no effect on insulin secretion nor islet apoptosis in the absence of pro-inflammatory cytokines. (A)** Static insulin secretion in response to basal (2mM) or 20mM glucose was assessed in isolated mouse islets incubated with either FK866 (10 nM) or compound 17 (200 nM) for 24 h. **(B)** Caspase 3/7 activity in mouse islets either untreated or exposed to FK866 (10 nM) for 24 h. N = 5 – 14 replicates from 6 mice and representative of three independent experiments.

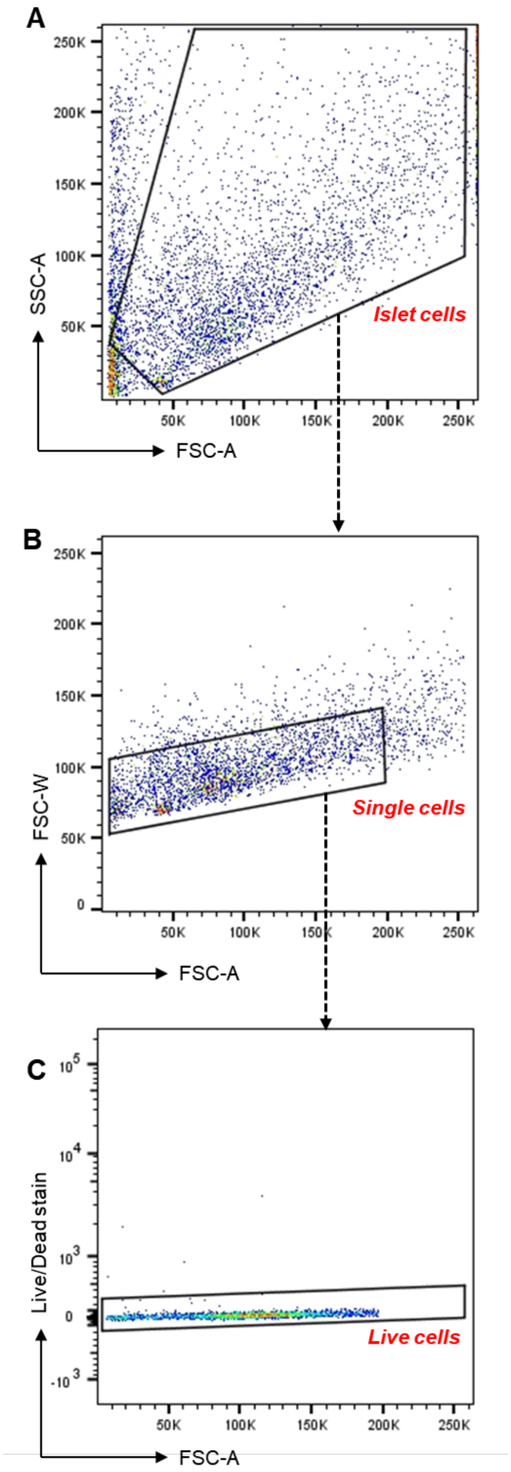


**Supplementary Fig. 3 Exemplary gating strategies for live isolated mouse pancreatic islet cells. Related to Figure 3, Figure 4, Figure 5 and S10**. Black polygons and gates are representative of (A) islet cell population, (B) islet single cell population (C) live cell population

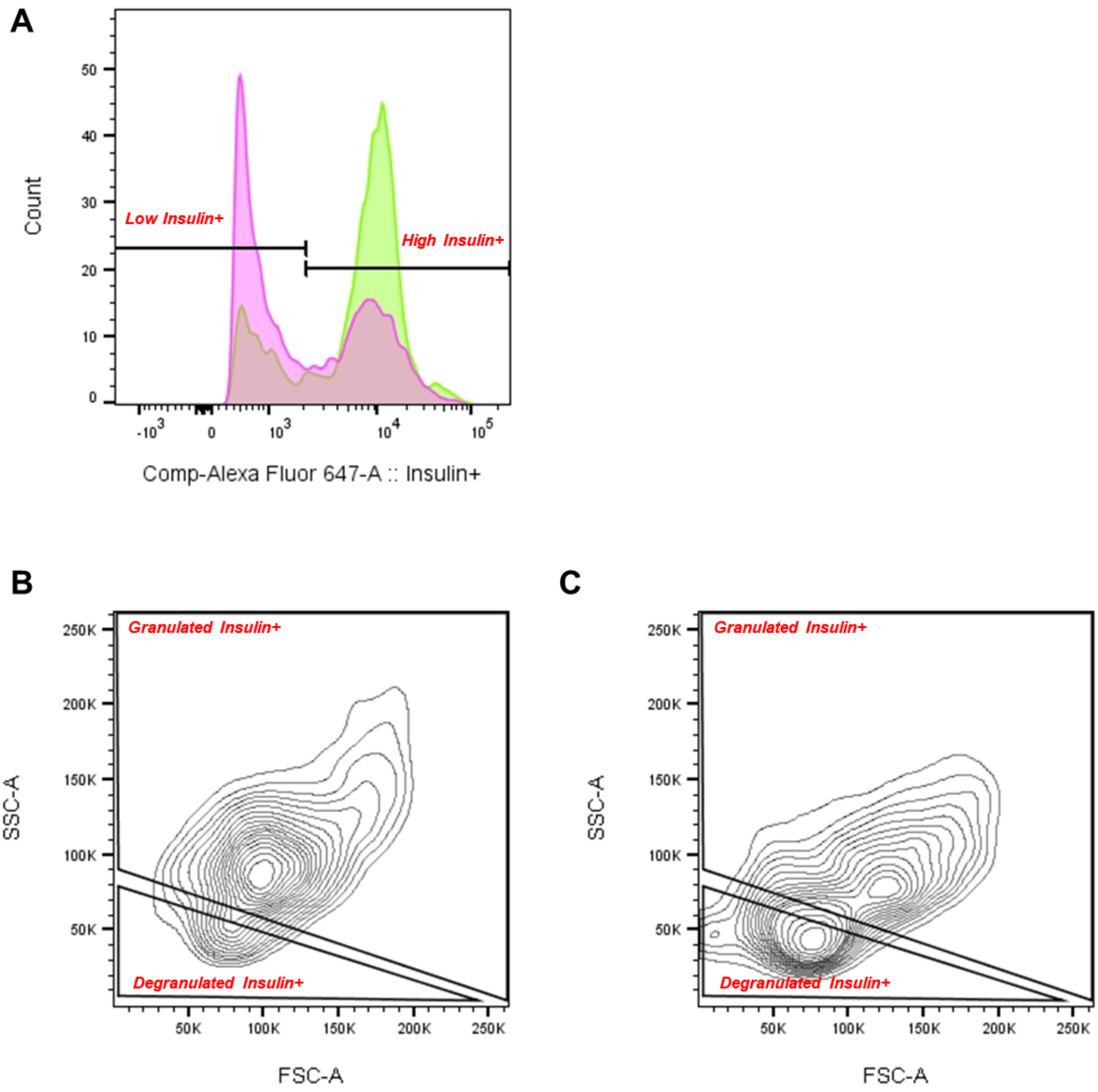


**Supplementary Fig. 4. Exemplary gating strategies for detecting granularity of isolated mouse pancreatic islet cells. Related to Figure 3.** Black polygons and gates are representative of (A) High and low Insulin + cells (B) Granulated and degranulated Insulin+ cells from non-diabetic control example. (C) Granulated and degranulated Insulin+ cells from diabetic control example. Green histograms represent non-diabetic control example. Pink histograms represent diabetic control example.

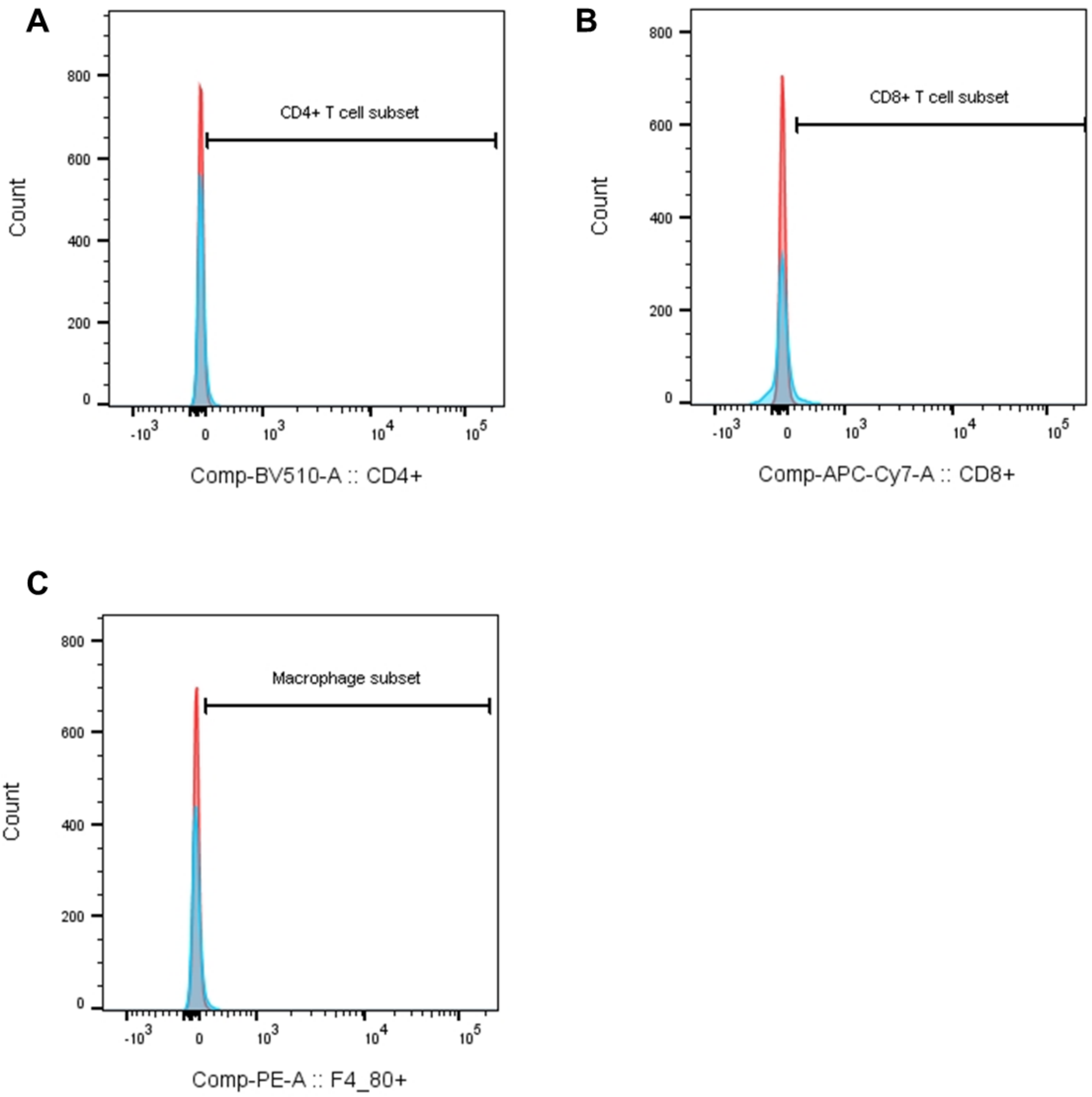


**Supplementary Fig. 5 Gating strategies for isolated mouse pancreatic islet cell immunofluorescent staining with immune cell markers. Related to Figure 4.** Black polygons and gates are representative of (A) CD4+ (B) CD8+ (C)F4/80+. Red histograms represented unstained cells. Blue histograms represent stained example population.

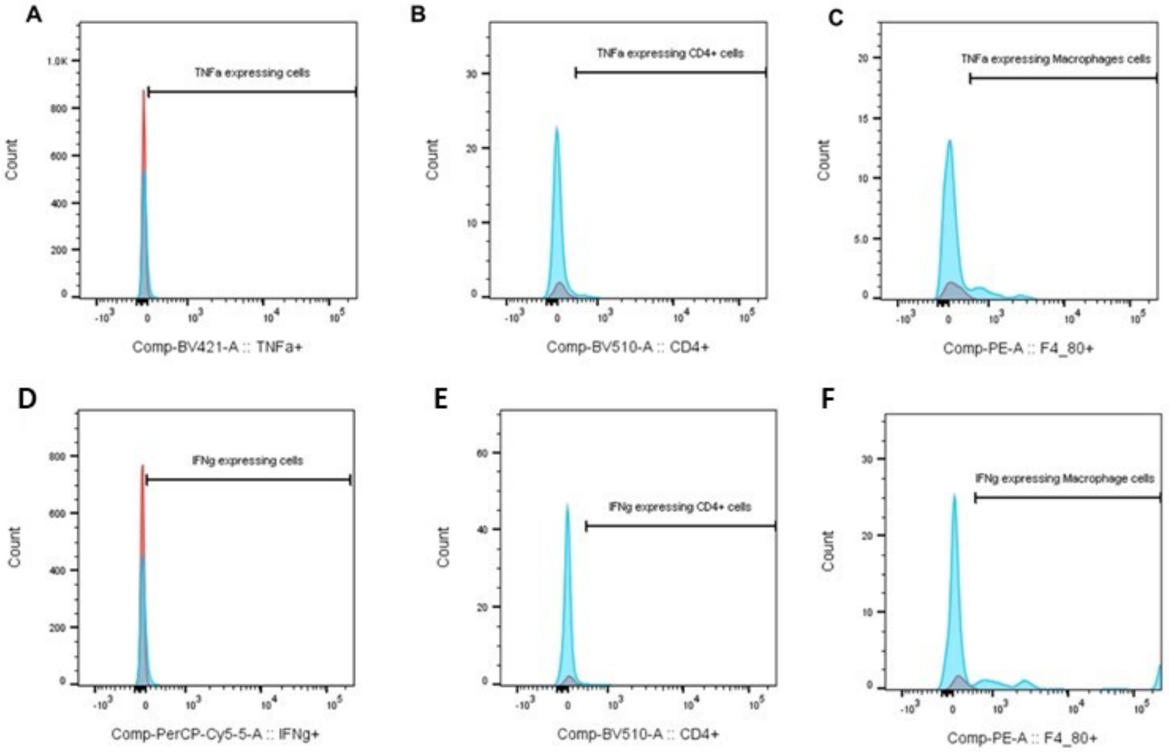


**Supplementary Fig. 6. Gating strategies for isolated mouse pancreatic islet cell immunofluorescent staining with inflammatory markers. Related to Figure 4 and Supplementary Fig. 10.** Black polygons and gates are representative of (A) TNFα+ (B) TNFα+ CD4+ (C) TNFα+ F4/80+ (D) IFNγ+ (E) IFNγ+ CD4+ (F) IFNγ+ F4/80+. Red histograms represented unstained cells. Blue histograms represent stained example population.

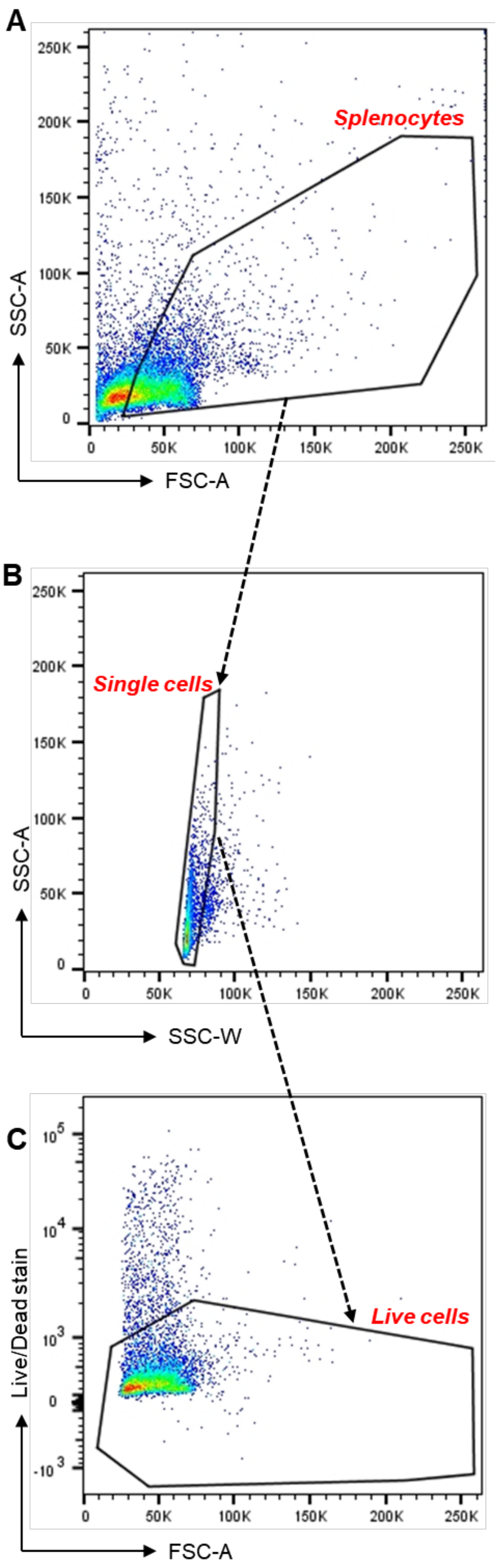


**Supplementary Fig. 7. Exemplary gating strategies for live isolated mouse splenocytes. Related to Figure 5**. Black polygons and gates are representative of (A) splenocyte population, (B) splenocyte single cell population (C) live cell population

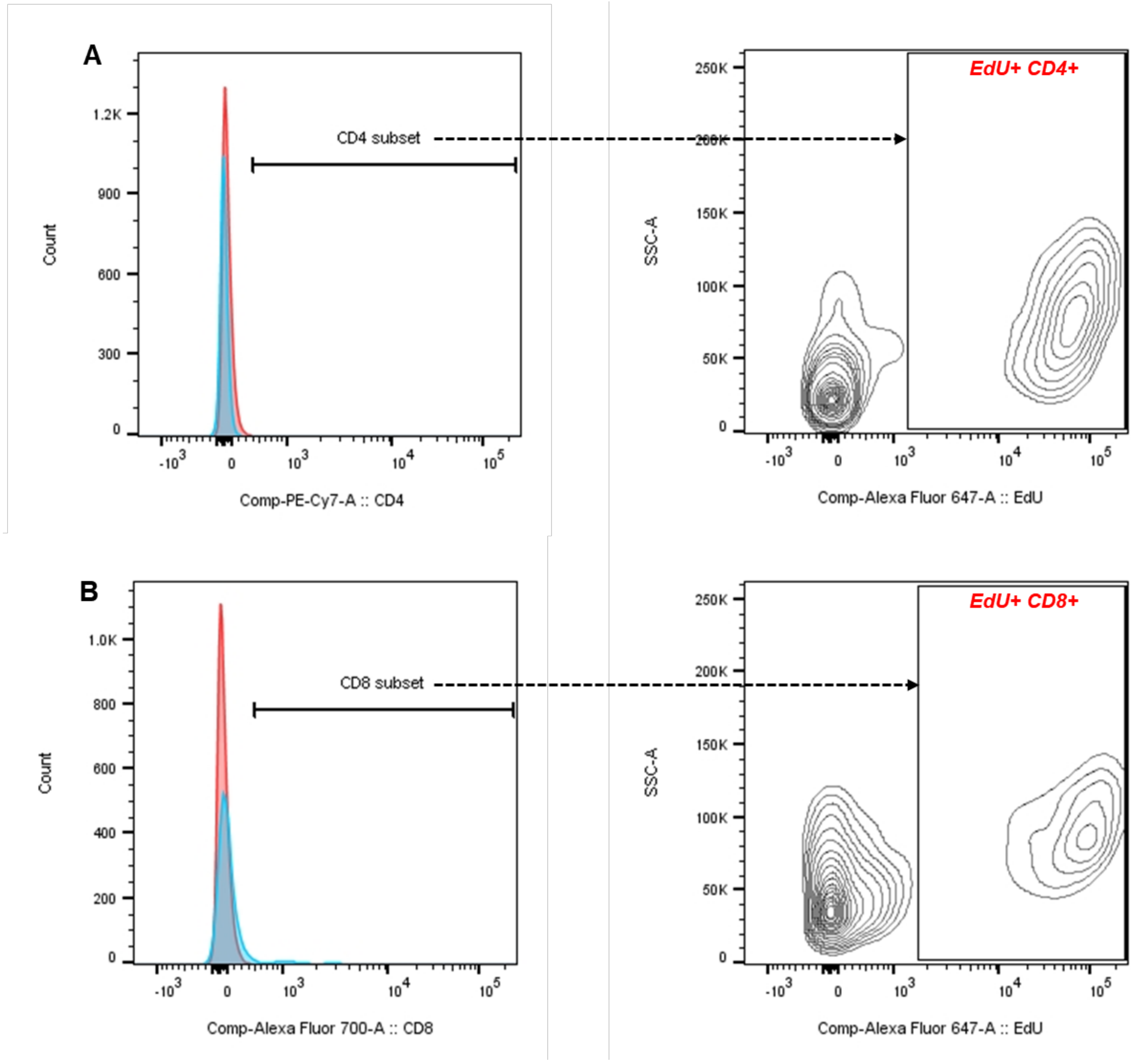


**Supplementary Fig. 8. Gating strategies for isolated mouse splenocyte immunofluorescent staining with immune cell markers and EdU+. Related to Figure 5.** Black polygons and gates are representative of (A) CD4+ and EdU+ CD4+ (B) CD8+ and EdU+ CD8+ cells. Red histograms represented unstained cells. Blue histograms represent stained example population.

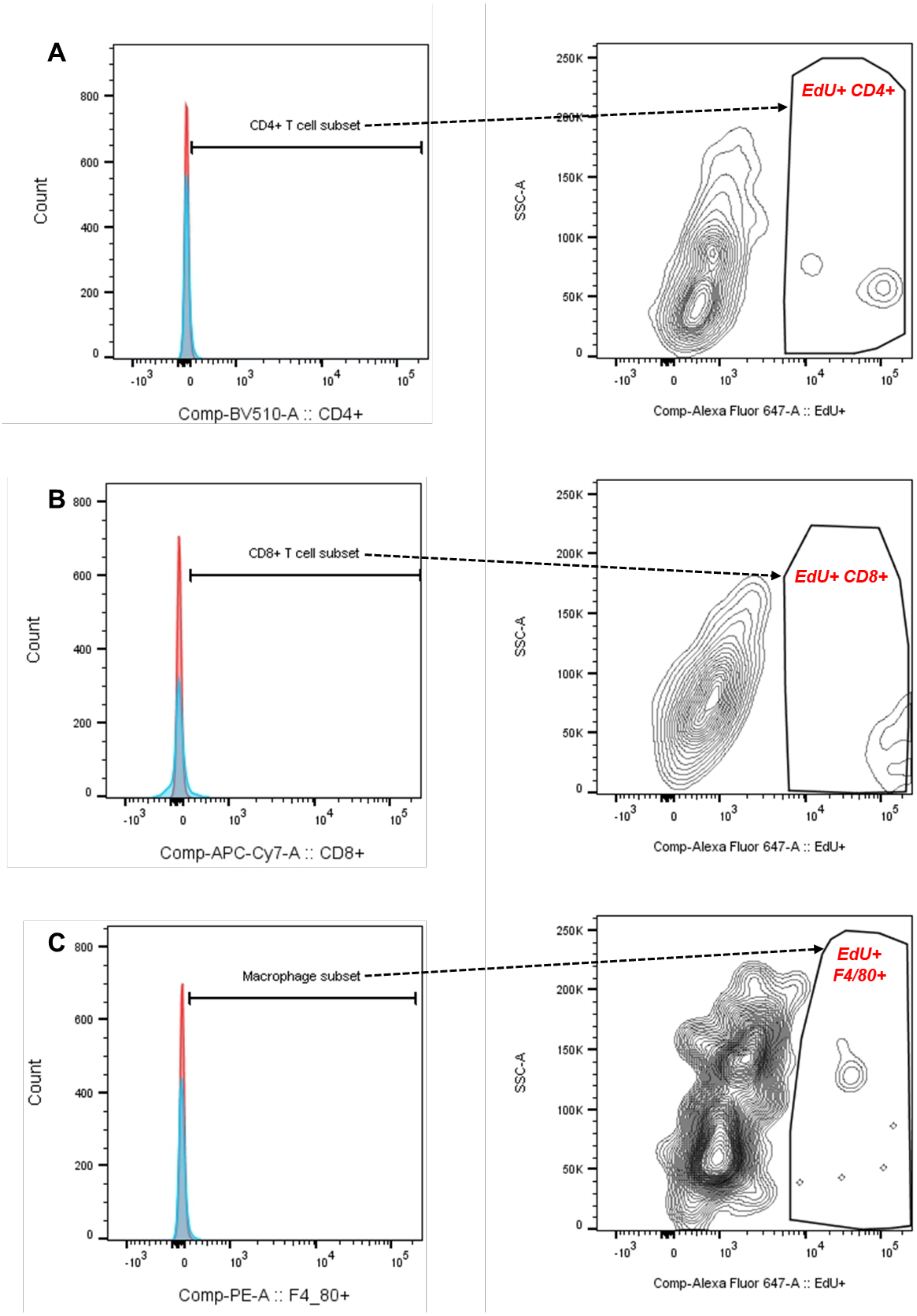


**Supplementary Fig. 9. Gating strategies for isolated mouse pancreatic islet cell immunofluorescent staining with immune cell markers and EdU+. Related to Figure 5.** Black polygons and gates are representative of (A) CD4+ and EdU+ CD4+ (B) CD8+ and EdU+ CD8+ (C) F4/80+ and EdU+ F4/80+ cells. Red histograms represented unstained cells. Blue histograms represent stained example population.

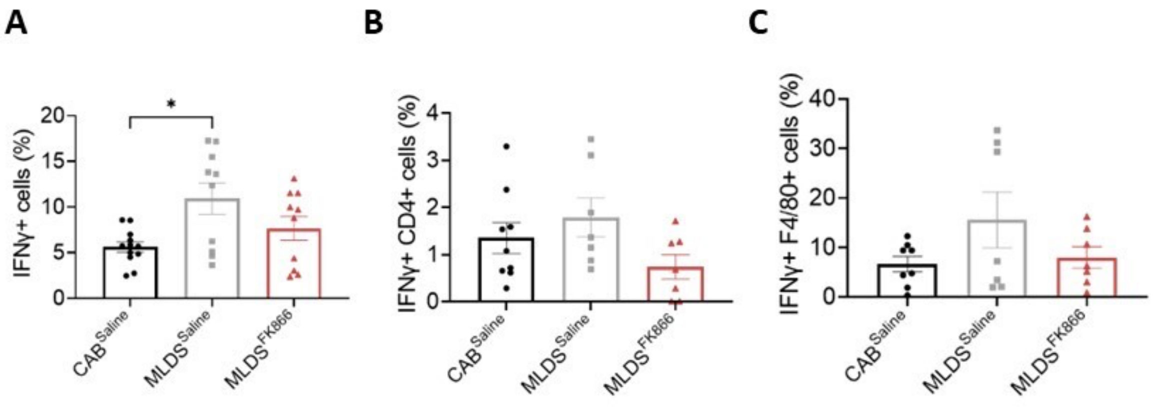


**Supplementary Fig. 10.** Percentages of **(A)** IFNγ+, **(B)** IFNγ+CD4+, **(C)** IFNγ+F4/80+, within the islet cell of CAB$^{Saline}$, MLDS$^{Saline}$ and MLDS$^{FK866}$ mice. n = 7-14 pools of islets of which a maximum of 5000 cells were analysed. Values in all graphs are represented as means ± SEM. Statistical significance is depicted by *p < 0.05.
